# Suppression rather than activation of the integrated-stress-response (GCN2-ATF4) pathway extends lifespan in the fly

**DOI:** 10.1101/2025.07.14.664701

**Authors:** Miriam S Götz, Dan J. Hayman, Gracie Adams, Fumiaki Obata, Mirre J P Simons

## Abstract

Stress response pathways are emerging as conserved modulators of lifespan. The prevailing hypothesis is that activation of stress responsive pathways, including the amino acid deprivation arm of the integrated stress response (ISR; the GCN2-ATF4 pathway) is pro-longevity. Activation of *ATF4* orthologs extends lifespan in *Saccharomyces cerevisiae* (yeast) and *Caenorhabditis elegans*, but its role in other longer-lived organisms remains unclear. We comprehensively tested the role of the GCN2-ATF4 pathway in the fly (*Drosophila melanogaster*) for the first time. We used conditional genetic manipulation of *dGCN2* and its downstream effector *Drosophila ATF4* (*crc; dATF4*). In contrast to previous studies, we show that overexpression of *dGCN2* and *dATF4* significantly reduces lifespan, while knockdown (in vivo RNAi) of *dATF4* extends lifespan. We confirmed *dATF4* activity was successfully modulated using a fluorescent *dATF4* activation reporter. Borrelidin, a tRNA synthetase inhibitor, significantly reduced lifespan in a both *dATF4* and diet-dependent manner, independent of microbial load, showing our modulation of *dATF4* altered nutrient to ISR signalling. We further conducted long-read RNA sequencing and found that our manipulation of *dATF4* changed global transcription in opposite directions, including known *ATF4* target genes. Enrichment analysis revealed that *dATF4* overexpression may drive metabolic stress, while *dATF4* knockdown upregulates proteostasis and DNA repair pathways. Our work reveals that *ATF4* exhibits a dual, dose- and context-dependent role in ageing. Chronic *dATF4* activation is detrimental in flies, while chronic suppression is pro-longevity. The GCN2-ATF4 pathway thus qualifies as a modifiable control of lifespan with cross-species relevance.

## Introduction

Stress response pathways are emerging as conserved modulators of lifespan (Haigis and Yankner, 2010; Kishimoto et al., 2018). Mild exposure to stress leads to increased stress resistance and lifespan extension through hormesis (Rattan et al., 2008). The activation of stress response pathways has further been suggested to be required for the longevity extension observed in many long lived mutants (most clearly demonstrated in *Caenorhabditis elegans*; Soo et al., 2023). Independently, overexpression of pathways that regulate stress responses (such as FOXOs, Nrf2, HIF; Hwangbo et al., 2004; Sykiotis and Bohmann, 2009; Bruns et al., 2015; Leiser and Kaeberlein, 2014; Rogers et al., 2023) and end-products of stress pathways such as heat-shock proteins (Tatar et al., 1997), may be sufficient to extend lifespan (Soo et al., 2023). The pathway that may be responsible for these shared stress responses, the Integrated Stress Response (ISR), a conserved stress-activated pathway, has been implicated in lifespan regulation across species (Derisbourg et al., 2021; Maragakis et al., 2023).

The ISR is activated by four kinases (General control non-depressible 2; GCN2, Protein Kinase R (PKR)-like endoplasmic reticulum kinase; PERK, PKR and Heme regulated inhibitor; HRI) in response to different types of stress (amino acid; AA) starvation, ER stress, viral infection and heme deprivation, respectively (Donnelly et al., 2013). Detection of stress (e.g., amino acid deficiencies via uncharged tRNAs and independently of tRNA via ribosomal arrest in the case of GCN2; Ye et al., 2010; Obata and Miura, 2024) by each of these kinases leads to phosphorylation of eukaryotic Initiation Factor 2 alpha (eIF2⍰; Ron, 2002). Phosphorylation of eIF2⍰ leads to a general reduction in global protein synthesis while at the same time selectively enhancing translation of Activating Transcription Factor 4 (ATF4) by enabling ribosomes to bypass inhibitory upstream Open Reading Frames (uORFs). However, mechanisms that activate ATF4 independent of eIF2⍰ phosphorylation although less well studied, but can be important (Kim et al., 2025). Surprisingly restriction of tyrosine, a non-essential amino acid, activates ATF4 independently of GCN2, yet the effects of tyrosine on lifespan are independent of ATF4 (Kosakamoto et al., 2022; Kosakamoto et al., 2024). Upon activation ATF4 acts as a key regulator of cell fate (Pakos-Zebrucka et al., 2016) by driving a broad transcriptional response thought to help cells adapt to stress (Harding et al., 2003). Importantly, ATF4’s target genes can either promote autophagy or apoptosis, depending on the intensity, duration and cellular context of the response (Neil and Masson, 2023).

Evidence from multiple species suggests that *ATF4* and its orthologs can affect lifespan. However, experimental evidence linking *ATF4* to lifespan is currently limited to its orthologs *GCN4* in *Saccharomyces cerevisiae* (yeast) and *atf-4* in *C. elegans*. Overexpression of *GCN4* and *atf-4* extends lifespan in *yeast* and *C. elegans*, respectively (Table 1; Mittal et al., 2017; Statzer et al., 2022), whereas knockdown reduces lifespan in yeast (Gulias et al., 2023) and had no effect on lifespan in *C. elegans* (Rousakis et al., 2013). Several more studies found that knockdown of *atf-4* removed lifespan extension conferred by other genetic or pharmacological interventions (Table 1). In vertebrates, *ATF4* has been indirectly linked to ageing and age-related phenotypes (Table 2), e.g., protection from muscle degeneration in the case of *ATF4* KO mice (Miller et al., 2023).

**Table 1.**
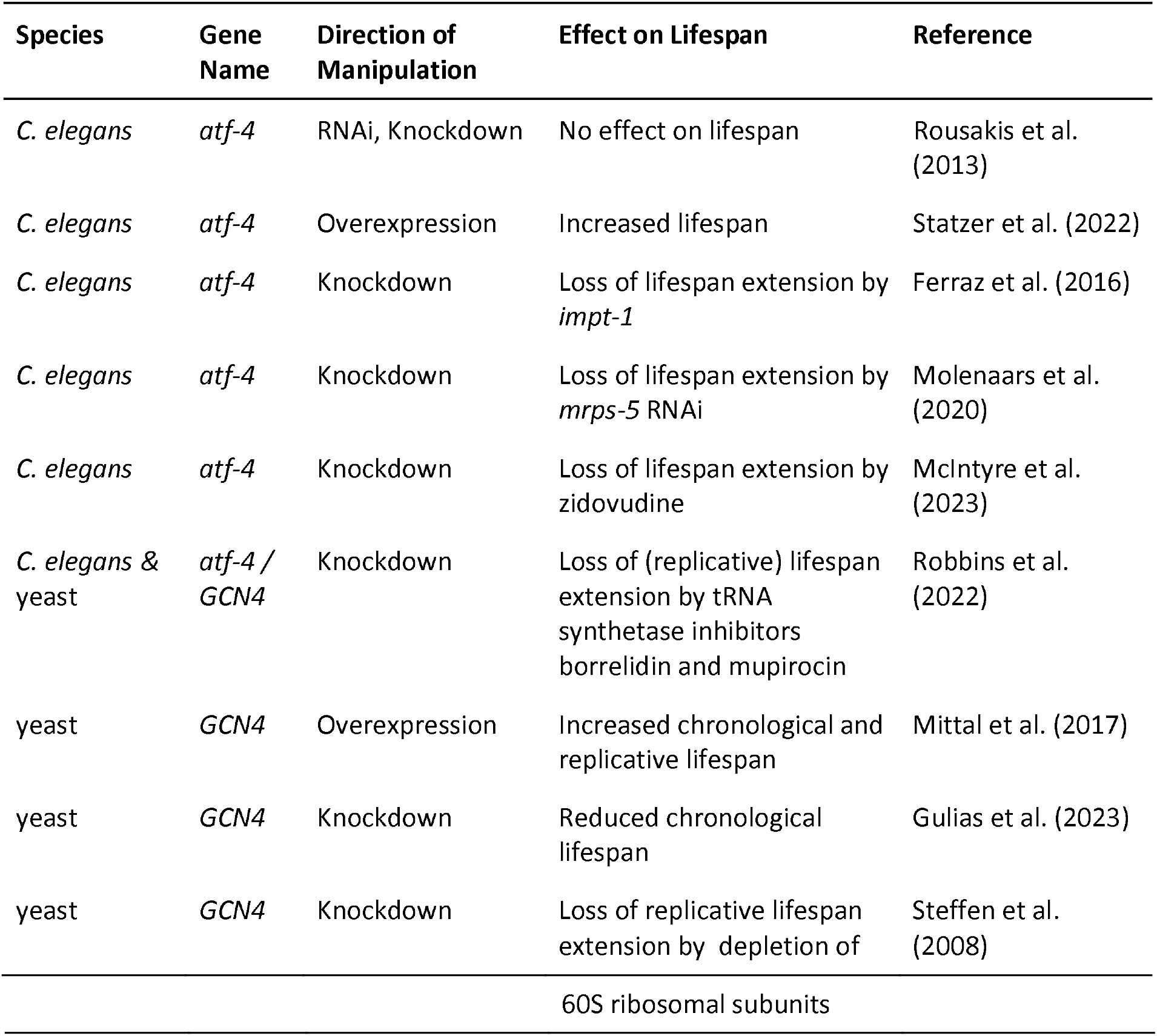
Lifespan effects of genetic manipulation of ATF4 orthologs. This table lists all published studies in which *ATF4* or one of its orthologs was genetically manipulated (e.g., overexpression or knockdown) and reported effects on whole-organism lifespan. Only yeast and C. elegans studies are available. Note *atf-4* is referred to as *atf-5* in literature published before 2022, *GCN4* is the yeast *ATF4* ortholog.

**Table 2.** ATF4 is linked to several age-related phenotypes across species. ↑Indicates upregulation of *ATF4*, ↓Indicates downregulation of *ATF4* either naturally, or as the result of genetic manipulation

| Species | *ATF4*<br>↑/↓ | Observed Phenotype | Reference |
| --- | --- | --- | --- |
| *M. musculus* & *D. melanogaster* | ↓ | *ATF4* knockdown alleviates diet-induced diabetes, hyperlipidemia and hepatosteatosis | Seo et al. (2009) |
| *M. musculus* | ↑ | Elevation of *ATF4* is a common feature in many long-lived mice mutants | Li et al. (2014)<br>Li and Miller (2015) |
| *M. musculus* | ↓ | Tissue specific knockdown of *ATF4* in the liver results in increased oxidative stress and increased free cholesterol | Fusakio et al. (2016) |
| *Danio rerio* | ↑ | Conditional *ATF4* overexpression promotes induces early onset of hyperlipidaemia and enhances adipogenesis in zebrafish | Yeh et al. (2017) |
| *M. musculus* | ↓ | Tissue specific knockdown of *ATF4* protects mice from common features of muscle ageing | Miller et al. (2023) |
| *M. musculus* | ↓ | *ATF4* drives regulatory T cell functional specification in homeostasis and obesity | Wang et al. (2025) |

In addition to genetic manipulation, pharmacological interventions have also been shown to manipulate the GCN2-ATF4 pathway (Piecyk et al., 2024). Notably, treatment with tRNA synthetase inhibitors has been shown to activate the GCN2–ATF4 pathway (Mariner et al., 2023; Carlson et al., 2023; Arita et al., 2017). Interestingly, treatment with borrelidin (threonine tRNA synthetase inhibitor) and mupirocin (isoleucyl tRNA synthetase inhibitor) has recently been suggested to extend lifespan in an *ATF4*-dependent manner in yeast and *C. elegans* (Robbins et al., 2022). Mechanistically, these inhibitors function by competing with their natural AA substrate, preventing proper charging of tRNAs. This process causes an accumulation of uncharged tRNAs, a hallmark of AA deprivation and a known trigger for GCN2 activation and subsequent upregulation of ATF4 translation (Krupitza and Thireos, 1990; Wek et al., 1989; Longchamp et al., 2018).

The direction of associations with *ATF4* is variable (Table 1 & 2). Still, the prevailing hypothesis, currently based on work in yeast and *C. elegans* and some associations in mice, combined with the idea that low stress activation is pro-longevity (Rattan, 2008), is that *ATF4* activation would lead to lifespan extension. There is however a lack of further experimental testing of the ISR pathway in ageing research, especially in physiologically more complex and longer lived organisms than yeast and *C. elegans*. In this study, we conditionally manipulated the ISR pathway in flies (*Drosophila melanogaster*) via both *dGCN2* and its downstream target, *cryptocephal* (crc; Drosophila ortholog of *ATF4*, hereafter referred to as *dATF4*) to determine their role in ageing. Experiments were conducted at fully fed and dietary restricted (DR) conditions, as mild stress and *ATF4* specifically have been implicated in the health benefits of DR (Jonsson et al., 2021; Rattan, 2008). We find contrary to the prevailing hypothesis that overexpression of *dATF4* shortened lifespan, whereas suppression of *dATF4* led to a robust lifespan extension.

## Results

### Suppression, not activation, of the GCN2–ATF4 pathway extends lifespan

To isolate effects during adult life, we used a conditional driver to overexpress or knock down *dGCN2* and its downstream effector *dATF4* during adulthood. A key benefit of using conditional drivers is that it fully controls for small genetic differences known to impact ageing and lifespan (Hayman et al., 2025; Tower, 2000). Ubiquitous overexpression of *dGCN2* during adult lifespan drastically reduced lifespan (*p* < 0.0001, *N* ≥ 354; Figure 1A; Table S1). *dGCN2* overexpression effects were significantly stronger on DR (*p* < 0.01, Table S1D). However, we interpret this as individuals dying rapidly irrespective of diet, with this effect appearing bigger on DR, as controls live longer on DR (*p* < 0.0001, Table S1C). In line with the *dGCN2* overexpression results, reduction in lifespan was also observed when *dATF4* was overexpressed (*p* < 0.0001, *N* ≥ 376; Figure 1C, Table S1A,B), independently of diet (p > 0.05, Table S1D). Knockdown of *dGCN2* resulted in a non-significant lifespan extension (*p* = 0.06, *N* ≥ 391; Figure 1C, Table S1A,B). Strikingly, when we knocked down *dATF4*, this significantly extended lifespan (*p* < 0.0001, *N* ≥ 363; Figure 1D, Table S2A,B) irrespective of diet (*p* > 0.05, Table S1D).

**Figure 1.**
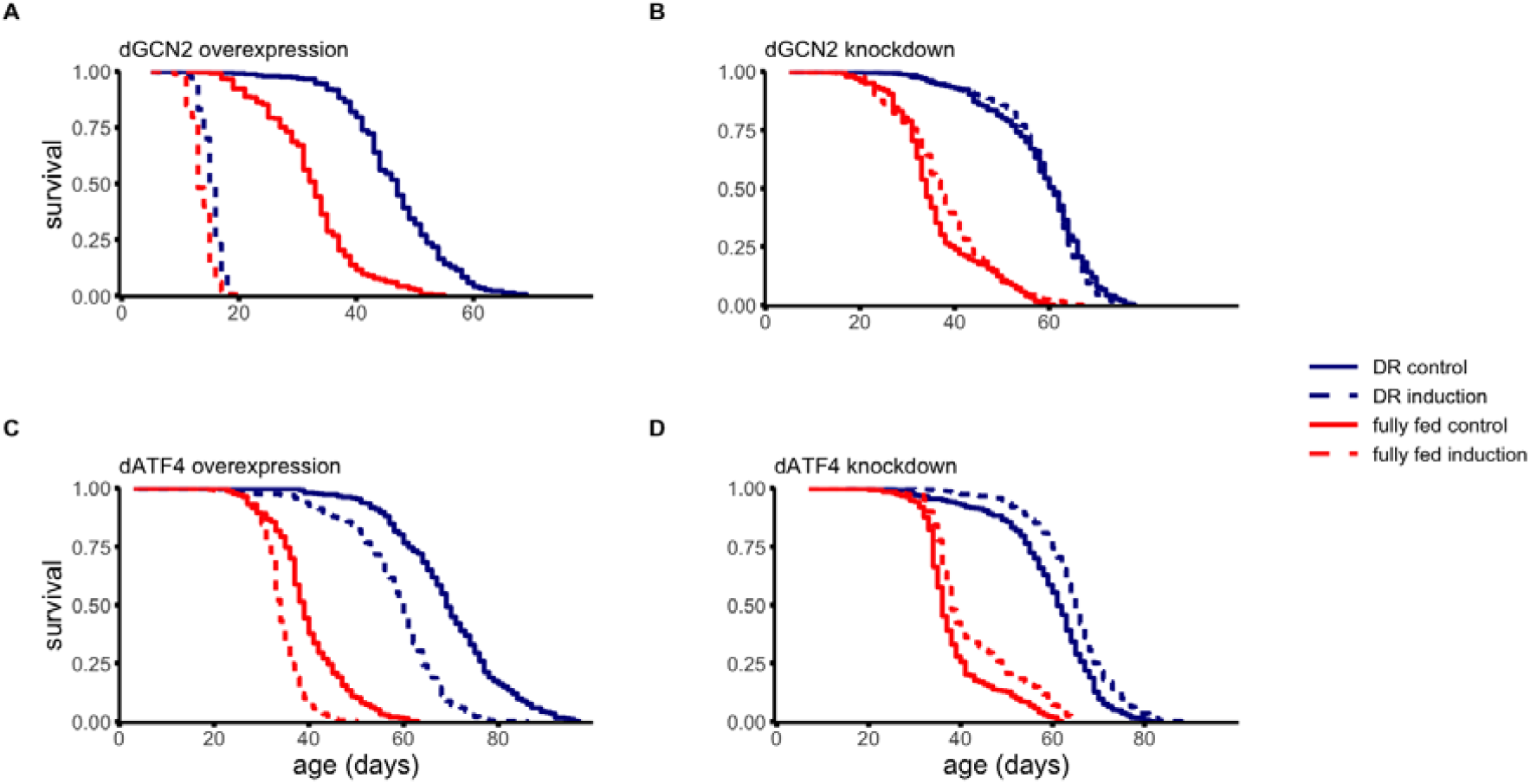
Lifespan is shortened by activation and extended by suppression of the GCN2-ATF4 pathway. Survival curves of flies with conditional induction of overexpression or knockdown (in vivo RNAi) of *dGCN2* (A,B) and *dATF4* (C,D). All sample sizes and statistics are given in Table S1.

### Conditional changes in dATF4 transcript levels correspond with reporter activity

As our results directly contradict prior findings on how *ATF4* modulates lifespan (Table 1), we wanted to confirm that in our experiments conditional changes in *dATF4* expression effectively altered downstream promoter activity. *ATF4* activity can be visualised using a fluorescent reporter, 4E-BP ^*intron*^-*dsRed* (Kang et al., 2017; Kosakamoto et al., 2022). The reporter contains an intron sequence with reported *dATF4* binding sites (Figure 2A). Conditional changes in *dATF4* expression were already visible 48h after conditional induction of knockdown and overexpression, respectively (Figure 2B,C). Induction of *dATF4* overexpression significantly increased reporter activity (*p* < 0.001, ANOVA, image analysis, Figure 2D; *p* < 0.05, Welch t-test, fluorescence spectroscopy, Figure 2F). A reduction in reporter activity was observed when *dATF4* was knocked down (Figure 2B, *p* < 0.01, ANOVA, microscopy, Figure 2D; *p* < 0.05, Welch t-test, plate reader, Figure 2E).

**Figure 2.**
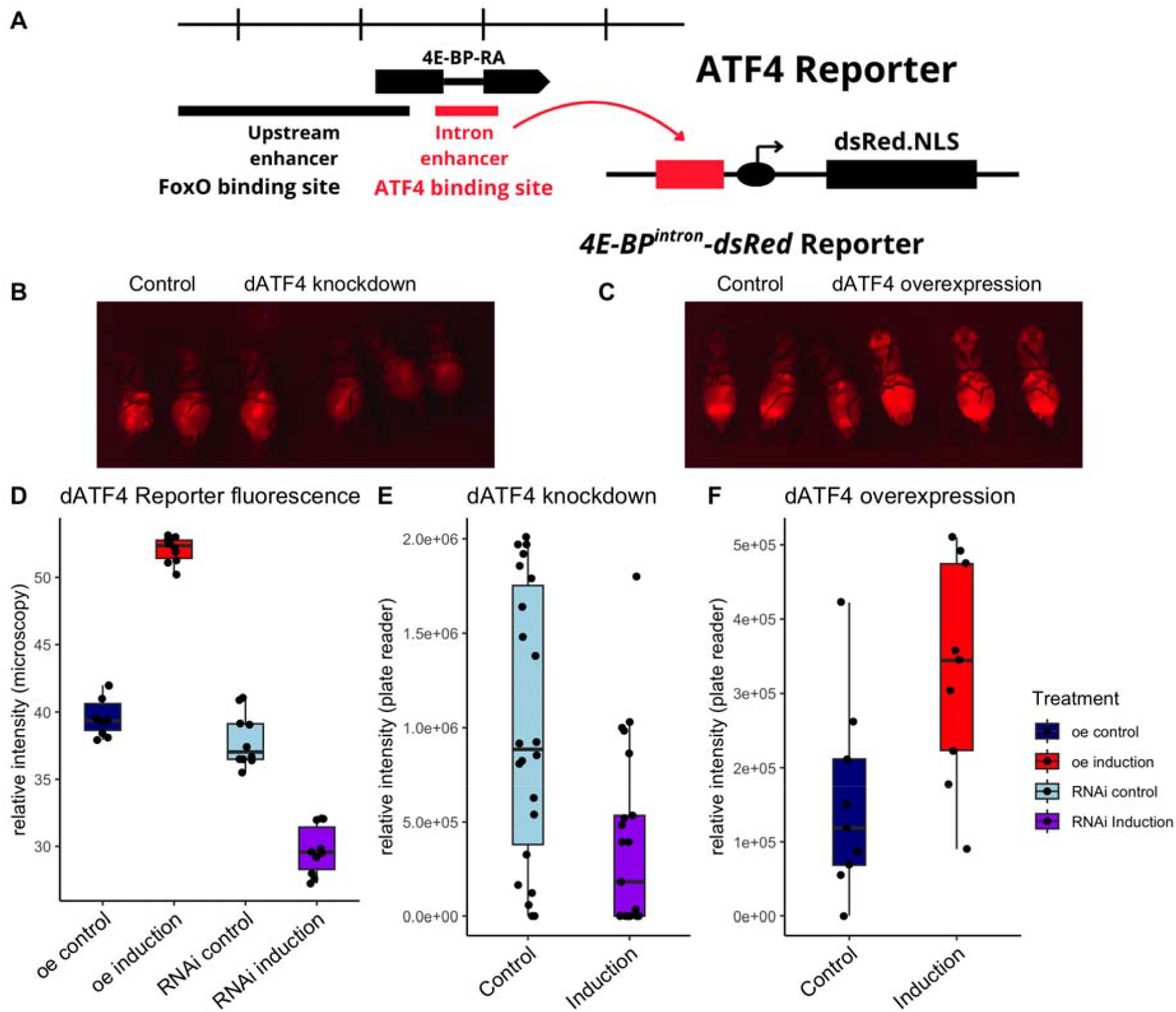
Conditional changes in *dATF4* transcript levels correspond with reporter activity. A, Structure of fluorescent *ATF4* reporter *4E-BP* ^*intron*^-*dsRed* (Adapted from Kosakamoto et al. (2022)). B,C, Representative images of conditional induction vs. control of (B) dATF4 overexpression and (C) *dATF4* knockdown. D,E,F Quantification of relative *4E-BP* ^*intron*^-*dsRed* fluorescence (D) in the whole fly from microscopy images using pixel intensity measurements in ImageJ and (E,F) from the body using fluorescent spectrophotometry. RFU resulting from (E) 50 flashes and (F) 10 flashes. B,C,

### Borrelidin modulates lifespan effects of dATF4 overexpression and knockdown

Pharmacological activation of the GCN2–ATF4 pathway, through tRNA synthetase inhibitors (specifically borrelidin), has recently been linked to lifespan extension in an *ATF4* and dose dependent manner (Robbins et al., 2022). In contrast to these findings, borrelidin reduced lifespan in flies (Figure 3, Table S2), in line with the lifespan reduction we observed when *dATF4* is overexpressed. Note, that dose is important here and we cannot exclude that at a lower dose or shorter exposure borrelidin can extend lifespan in the fly. Intriguingly borrelidin did not decrease lifespan further when *dATF4* was overexpressed (Figure 3A, Table S2C) and removed the lifespan extending effect of *dATF4* suppression (Figure 3B, Table S2D). The interaction between borrelidin and genetic manipulation of *dATF4* expression levels in our experiments does indicate our genetic modification modulates the ISR response effectively.

**Figure 3.**
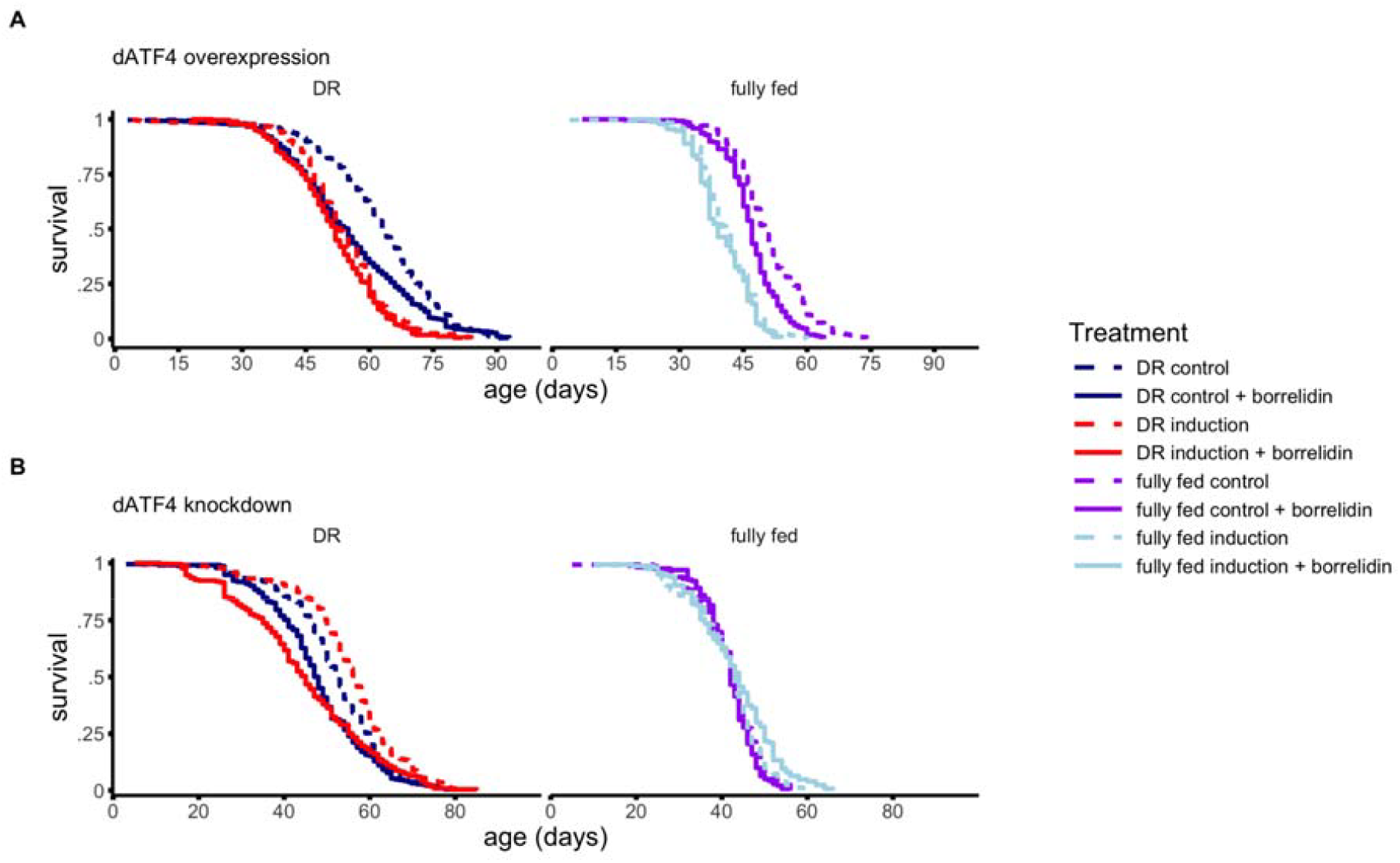
Borrelidin modulates the lifespan effects of *dATF4* and reduces lifespan (A, B). (A) Borrelidin did not reduce lifespan when *dATF4* was overexpressed. (B) Conversely, *dATF4* knockdown failed to extend lifespan when flies were treated with borrelidin. All sample sizes and statistics in Table S2. Note sample sizes for fully fed conditions are lower especially for the *dATF4* knockdown condition (B, right panel), and these results should therefore be considered less informative.

### Axenic conditions do not interact with the lifespan modulation of dATF4

The described biological actions of borrelidin include antimicrobial effects (Fang et al., 2015). In addition, *ATF4* has been implicated in immunity (Wang et al., 2025; Mukherjee et al., 2020). To confirm that any observed changes in lifespan following borrelidin could be attributed to tRNA synthetase inhibition rather than its antibiotic properties, we treated conditional *dATF4* overexpression and knockdown flies with a broad range antibiotic to eliminate their microbiome. Antibiotic treatment had no significant effect on lifespan in either *dATF4* overexpression or knockdown flies (p > 0.05, Figure S1, Table S3, respectively) and did not modulate lifespan changes observed via genetic manipulation of *dATF4* (*p* > 0.05, Figure S1, Table S4). These results suggest that the observed lifespan changes are driven by borrelidin’s effect on the GCN2–ATF4 pathway, rather than its antibiotic properties or effects of *ATF4* on immunity which would likely lead to a differential response in axenic conditions.

### Transcriptomic response to dATF4 overexpression and knockdown

To explore the molecular mechanisms underlying the observed lifespan extension and reduction following conditional *dATF4* manipulation we conducted ONT Transcriptome Sequencing. We measured the transcriptional response of whole flies 10 days after induction of dATF4 overexpression or knockdown in comparison to their respective controls. Similar to lifespan experiments, flies were kept either on fully fed or DR conditions. Given the lack of consistent diet-by-genotype interactions in lifespan (Figure S2, Table S1D, Table S4), we focused on transcriptomic differences driven by transgene induction alone whilst controlling for diet. We observed a modest, but statistically significant negative correlation in differential gene expression in response to *dATF4* knockdown and *dATF4* overexpression, suggesting *ATF4* targets are effectively modulated in our experiments (Figure 4). Furthermore, when we selected genes that changed significantly in expression (p < 0.01, not FDR corrected) we found 18 genes that changed in both conditions, and of these 13 changed in the opposite directions. Although this overlap is not larger than expected by chance (*p* > 0.05) several of these genes have been previously reported to respond to *ATF4* across species, and their observed directional change in transcription matches with the predicted response in the literature (Table S5), most notably *MAT2A* and *SLC36A1*. As such the analysis of significant gene overlap appears to support the biological conclusion that our experimental manipulation of ATF4 levels changes the transcriptome in an *ATF4* dependent manner as intended.

**Fig. 4.**
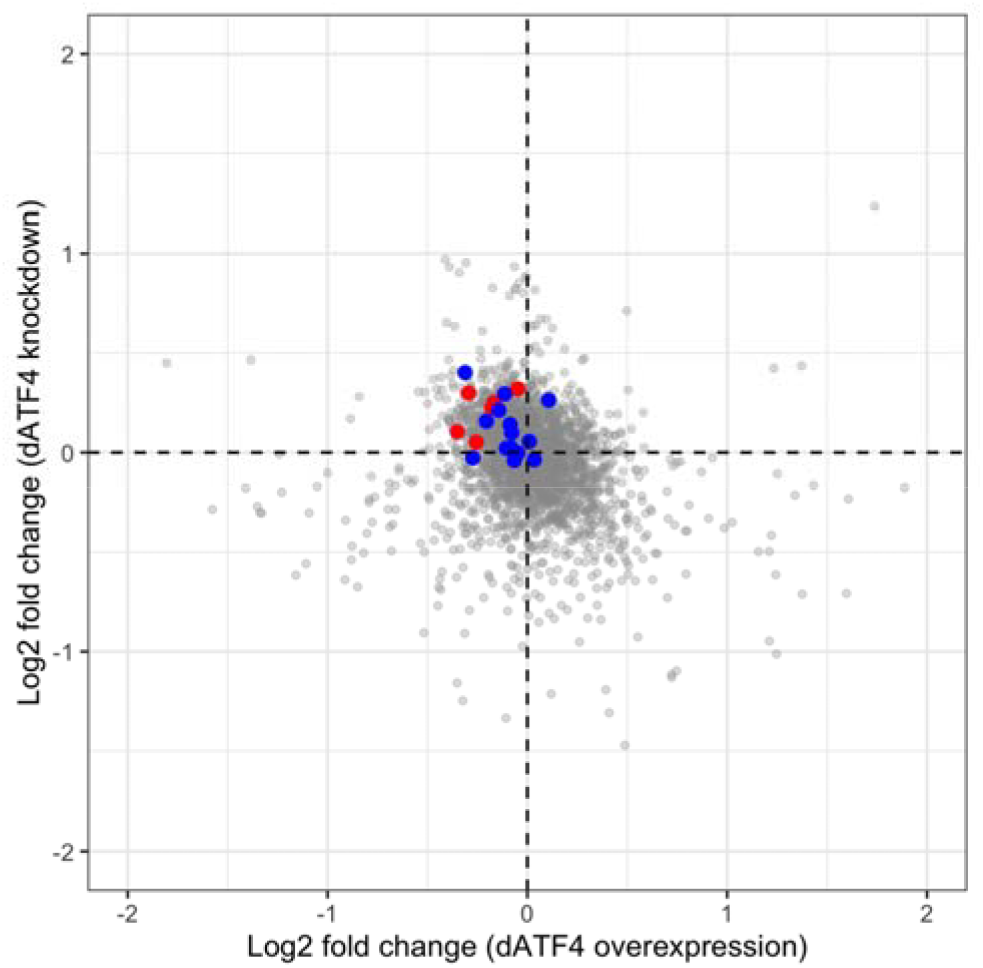
Rank correlation (−0.28, *p* < 0.0001) between differential expression in response to *dATF4* overexpression and knockdown suggesting concordant transcriptional change in response to *dATF4* directional manipulation. Highlighted genes from two concordant enrichment categories are highlighted, DNA mismatch repair (red) and Nucleotide excision repair (blue).

KEGG-based gene set enrichment analysis was used to better understand which molecular pathways were significantly enriched in *dATF4* overexpression and knockdown. We show the top 10 enriched up and down regulated terms irrespective of false discovery rate. This revealed distinct transcriptomic profiles for *dATF4* overexpression and knockdown respectively, however interestingly we found that two DNA repair pathways (Mismatch repair and Nucleotide excision repair) were enriched in opposite directions, i.e. enriched in *dATF4* knockdown and suppressed in dATF4 overexpression (Figure 4).

KEGG pathway analysis of *dATF4* overexpression (Figure 5A) reveals downregulation of DNA replication and mismatch repair and nucleotide excision repair, possibly indicating a breakdown of cellular maintenance. Furthermore, several metabolic pathways, including starch and sucrose metabolism and galactose metabolism are downregulated, suggesting a metabolic shift away from carbohydrate metabolism. Simultaneously we observe an upregulation of glutathione metabolism, a pathway involved in detoxification processes, often associated with oxidative stress, and aminoacyl-tRNA biosynthesis and Glycine, serine and threonine metabolism, possibly reflecting stress-related changes in protein homeostasis. Intriguingly, further GO term analysis suggests reduced capacity to respond to stress (Figure 5B).

**Figure 5.**
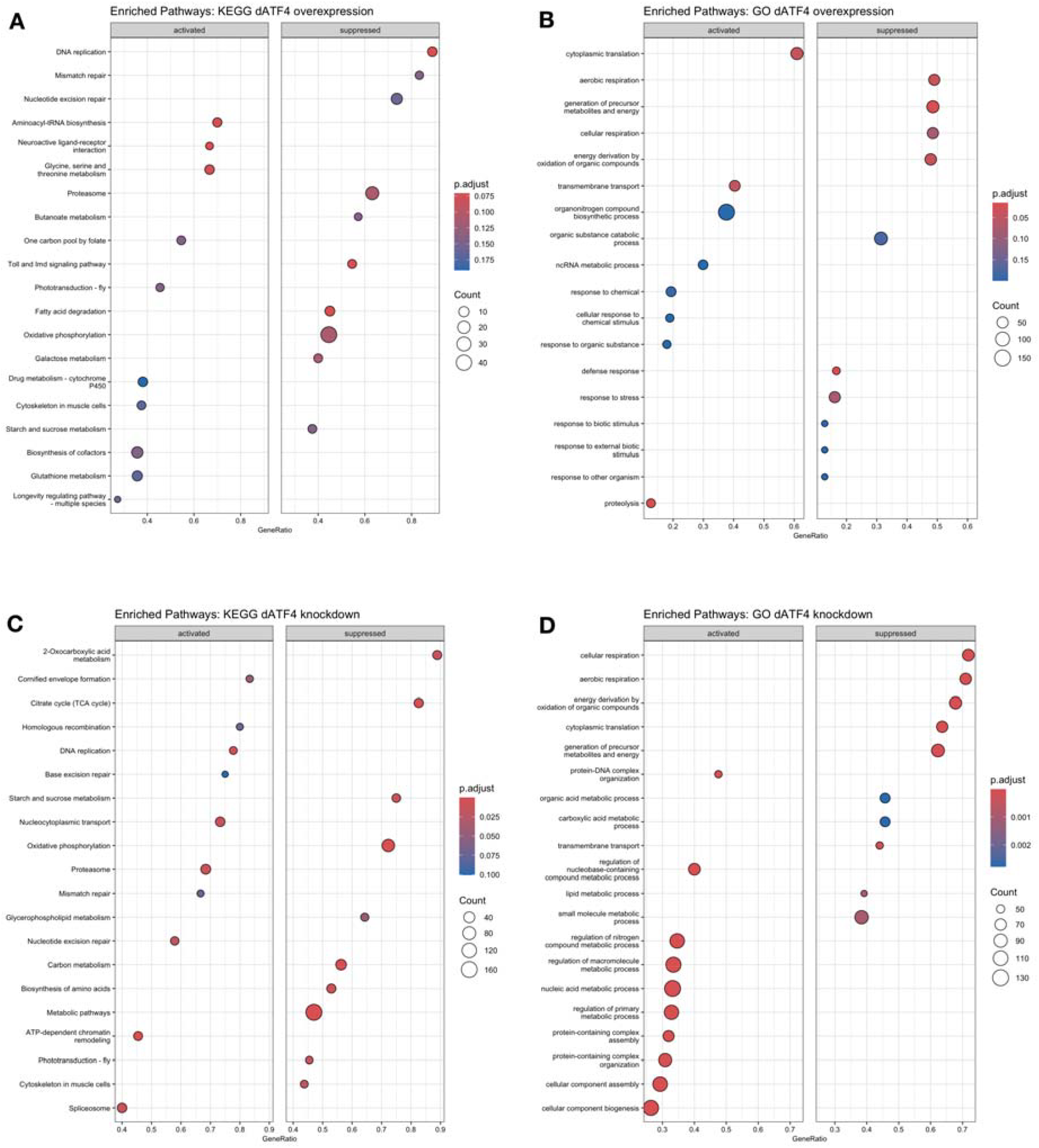
Top 10 activated and suppressed KEGG Gene Set Enrichment Analysis and GO terms, respectively for the effect of induction of *dATF4* overexpression and knockdown The data are split for pathways that were activated or suppressed in (A,B) *dATF4* overexpression and (C,D) *dATF4* knockdown (*in vivo* RNAi)

Although *dATF4* knockdown resulted in a smaller number of DEGs, KEGG pathway analysis revealed distinct transcriptional shifts, particularly in pathways associated with energy metabolism, protein synthesis and cellular maintenance (Figure 5C). Oxidative phosphorylation and the citrate cycle (TCA cycle) are downregulated, which suggests reduced energy expenditure. Alongside this, AA biosynthesis and ribosome-related pathways were also downregulated, suggesting a broader suppression of anabolic activity and translation. In combination, these changes may reflect a shift away from energy-expensive processes and a reduction in cellular workload. Upregulation of DNA repair pathways may reflect either a response to increased damage or a shift toward preserving genome integrity, resulting in a healthier physiological state. Furthermore, KEGG shows upregulation of the proteasome pathway and further GO analysis shows upregulation of GO terms associated with increased quality control (e.g., protein containing complex assembly, RNA metabolic process, and cellular component assembly; Figure 5D). This suggests that loss of *dATF4* may trigger compensatory mechanisms to maintain cellular homeostasis and this may lead to the observed lifespan extension.

### dATF4 lifespan effect is consistent across three replicates

To analyse the consistency of *dATF4* manipulation on lifespan we compared the primary *dATF4* lifespan results (Figure 1) to a re-analysis of the control groups of two independent experiments (Figure 3 & S1). These replicates were conducted in the same lab, but at different times and differ only in sample size and vehicle control added to the food (see methods). Overexpression of *dATF4* reduced lifespan across all experiments (Table S4C). In one experiment, the effect of overexpression on lifespan was smaller on DR. Similarly but in the opposite direction, *dATF4* knockdown extended lifespan in all experiments. In one experiment effects are again weaker in DR, but at this moment we have no strong evidence that the effects of *ATF4* on lifespan are diet-dependent. The effects of *dATF4* on lifespan are remarkably similar, making us confident that its effect on lifespan we report is robust.

## Discussion

Overexpression of *ATF4* orthologs in yeast and *C. elegans* has been associated with lifespan extension. In contrast, our work shows that activation of the GCN2–ATF4 pathway, via borrelidin, overexpression of *dGCN2* and its downstream effector *dATF4*, significantly reduces lifespan. Importantly ATF4 remains an important geroscience target as *dATF4* knockdown resulted in a robust lifespan increase. Long-read RNA-seq experiments suggest that under induction of *dATF4* overexpression, flies show transcriptional signatures that may be consistent with a shift away from carbohydrate-based energy metabolism and upregulation of detoxification pathways (e.g., glutathione metabolism) which may be consistent with increased stress, while *dATF4* knockdown may lead to increased investment in cellular maintenance via upregulation of proteostasis and DNA repair. These findings highlight the GCN2–ATF4 pathway as a modifiable regulator of lifespan in flies, with potential cross-species relevance.

Exposure to mild stress and the cellular stress responses they activate are generally thought to extend lifespan (Cypser and Johnson, 2002; Dues et al., 2016; Heidler et al., 2010; Schieber and Chandel, 2014; Soo et al., 2023). Overexpression of stress responsive transcription factors is generally associated with increased longevity, most clearly and repeatedly demonstrated in C. elegans (Senchuk et al., 2018; Hsu et al., 2003; Sural et al., 2019; Tullet et al., 2008; Zhang et al., 2009). ATF4 has been proposed to play a similar pro-longevity role (Table 1). However, our work shows that conditional overexpression of dATF4 reduces lifespan in flies, challenging the prevailing interpretation that ATF4 is universally beneficial. Indeed, other studies, again mainly in C. elegans suggest that the relationship between stress pathway activation and lifespan is context-dependent, and may also depend on the tissue or timing of expression. For example, constitutive overexpression of ATFS-1 has been shown to reduce lifespan (Bennett et al., 2014; Soo and Van Raamsdonk, 2021) and constitutive overexpression of XBP-1 has tissue-specific effects on lifespan. Ubiquitous and muscle-specific overexpression of XBP-1 reduces lifespan, while tissue-specific overexpression in the neurones or the intestine extends lifespan (Taylor et al., 2013). Furthermore, the reported lifespan effects of the ISR are variable, with manipulation of some pathway genes (e.g., *eIF2Bγ/ppp-1* and *eIF2αS51A*; Derisbourg et al., 2020) suggesting that suppression of the ISR promotes longevity (Derisbourg et al., 2021). Our results add to this evidence by demonstrating that sustained systemic activation of the GCN2-ATF4 pathway can have deleterious effects, while suppression of the pathway extends lifespan.

One explanation for the contradictory findings across studies may lie in the complex and context-dependent nature of *ATF4* signalling. Low level or short term activation of *ATF4* leads to upregulation of pro-survival pathways (Harding et al., 2003; Novoa et al., 2003); *ATF4* activation at high levels or for an extended period of time may result in upregulation of pro-apoptotic pathways (Frank et al., 2010; Han et al., 2013; Hiramatsu et al., 2014; Ishizawa et al., 2019; Pakos-Zebrucka et al., 2016). Our findings suggest a more nuanced picture of the role of ATF4 in the biology of ageing and challenge the idea that more *ATF4* equates to an improved physiological state (Mittal et al., 2017, Statzer et al., 2022; Robbins et al., 2022; Mariner et al., 2024). This supports a model in which chronic or intense *ATF4* activation leads to maladaptive effects and aligns with a growing body of evidence that *ATF4* activation effects are multifaceted (Seo et al., 2009; Wang et al., 2025; Miyake et al., 2016; Adams et al., 2017). For example, ATF4 knockout mice are protected from age-related muscle degeneration (Miller et al., 2023). Furthermore, *ATF4* activation is frequently hijacked by tumour cells to enhance cell survival from the stress that results from rapid proliferation and nutrient limitation (Hao et al., 2016; Ameri et al., 2004). However, increased *ATF4* activity was also found to sensitise tumour cells to therapy-induced cell death (Ishizawa et al., 2016; Zong et al., 2017). This highlights that *ATF4*-driven stress responses may confer either resilience or vulnerability, depending on the context (Wortel et al., 2017).

Interestingly, two types of tRNA synthetase inhibitors, including borrelidin, are thought to extend lifespan in yeast and *C. elegans* in an *ATF4*- and dose-dependent manner (Table 1). Additionally, other work in both *C. elegans* (Lee et al., 2003; Webster et al., 2017) and flies (Suh et al., 2020) has shown that RNAi of tRNA synthetases is associated with lifespan extension. Borrelidin, a known tRNA synthetase inhibitor, activates the GCN2-ATF4 pathway via accumulation of uncharged tRNAs, thereby mimicking AA deprivation (Francklyn and Mullen, 2019). This positions borrelidin as a mechanistic probe of the GCN2-ATF4 pathway. However, we find that, at least at our tested dose, borrelidin significantly reduces lifespan. Consistent with our findings that overexpression of *dGCN2* and its downstream target *dATF4* reduces lifespan, we find that pharmacological activation of this pathway using borrelidin reduces lifespan in an *dATF4* dependent manner.

We find that borrelidin does not reduce lifespan when *dATF4* is overexpressed and that borrelidin removes the lifespan extension resulting from *dATF4* knockdown. Perhaps overexpression of *dATF4* is already exerting its maximum negative effect on lifespan and additional pharmacological activation by borrelidin is therefore not observed. However, in this scenario we would expect knockdown of *dATF4* to rescue the negative effects activation of *dATF4* by borrelidin has on lifespan. This is not the case, as on DR, knockdown of *dATF4* exacerbates the lifespan shortening by borrelidin, suggesting more complex regulation of *ATF4s* targets other than through *ATF4* abundance.

A possible explanation for these results is that the experimental modulation of *dATF4* transcript levels changes posttranslational modifications of *ATF4*. The relative posttranslational state of *ATF4* is understood to be complex and the reason why *ATF4* activation can both lead to suppression and activation of its target genes (Neil and Masson, 2023; Wortel et al., 2017; Zhang et al., 2024). Borrelidin may independently modulate said posttranslational modifications through GCN2 and may therefore not lead to the expected directional relationships. As such our wider results could indicate that *dATF4* knockdown extends lifespan as it leads to a more pro-longevity genomic transcriptional signature, perhaps not through the abundance of *dATF4* protein, but its relative posttranslational state. This is an intriguing yet currently speculative hypothesis that will require further testing.

We hypothesised that the effect of *ATF4* on lifespan is dependent on both intensity and duration of activation, with chronic or strong activation potentially inducing more apoptosis related target genes (Neil and Masson, 2023). Our long-read RNA-seq results revealed transcriptional signatures indicative of a shift away from carbohydrate-based energy metabolism and reduction in DNA repair mechanisms. Simultaneously flies upregulated detoxification pathways, including glutathione metabolism, which is important for responding to oxidative stress (Wilhelm et al., 1997; Bell et al., 2011; Kreß et al., 2023). The observed upregulation of aminoacyl-tRNA biosynthesis is consistent with findings that suggest tRNA synthetases may reflect a compensatory mechanism that is activated under chronic activation of the ISR (Jones et al., 2023) and could explain why further activation of the GCN2-ATF4 pathway via tRNA synthetase inhibition had limited additional effects on lifespan in *dATF4*-overexpressing flies. In combination, this transcriptional profile suggests a downregulation of key reproductive and metabolic processes potentially reflecting a stress-induced reprioritisation of transcriptional programmes (Weiße et al., 2015; Kim et al., 2020; Kim et al., 2021). Together, these data suggest that chronic *dATF4* activation induces a metabolically costly transcriptional state that deprioritises reproduction, increases stress resistance pathways that may prove hard to sustain under chronic activation of *dATF4*, ultimately increasing physiological stress or predisposing cells to apoptosis (Meng et al., 2019; Wortel et al., 2017; Mukherjee et al, 2020).

In contrast to the stress-associated signature of the overexpression data, we find that dATF4 knockdown resulted in a downregulation of oxidative phosphorylation and the TCA cycle. This contrasts with previous findings in pro-longevity contexts, which link upregulation of oxidative phosphorylation to lifespan extension in DR and long-lived mutant mice (Gautrey et al., 2025; Elmansi and Miller, 2023). Notably, in a cortical neuron model, *ATF4* is thought to be a pro-death transcription factor with *ATF4* knockdown in cortical neurons thought to increase resistance to oxidative death (Lange et al., 2008). However, the overall suppression of anabolic pathways (including AA biosynthesis and ribosome-related pathways) is consistent with other pro longevity phenotypes (Hou et al., 2016). Furthermore, we found various DNA repair pathways enriched in *dATF4* knockdown flies. Although this may reflect a response to increased damage, it could also be a sign of increased cellular maintenance (McCracken, Adams et al., 2020). Additionally, we observe upregulation of proteasome and ubiquitin-mediated proteolysis pathways, which contribute to cellular maintenance. Previous work on *C. elegans* suggests that upregulation of the proteasome in response to mitochondrial stress, potentially caused by a downregulation of oxidative phosphorylation, leads to lifespan extension (Sladowska et al., 2021). Together, these findings suggest that the pro-longevity effect of *dATF4* knockdown may be the result of a coordinated shift toward cellular maintenance. This pattern may reduce metabolic strain, helping to extend lifespan in *dATF4* knockdown flies. Understanding the precise downstream targets of *ATF4* that modulate lifespan and how *ATF4* regulates its transcription will help understand where the reported anti-ageing effects of the ISR originate.

Our study challenges the prevailing hypothesis that ATF4 activation is broadly beneficial for lifespan by demonstrating that, in flies, chronic or systemic activation of the GCN2-ATF4 pathway is detrimental, while suppression of *dATF4* extends lifespan. These findings suggest that the role of *ATF4* in ageing is likely to be dose-, and context-dependent. Our work now further positions the GCN2-ATF4 pathway as a modifiable regulator of lifespan, with potential relevance across species.

## Methods

### Fly husbandry and diets

All flies were reared on “fully fed” fly media (8% yeast), prepared to the following specifications: 8% yeast, 13% table sugar, 6% cornmeal, 1% agar and 0.225% nipagin (all w/v) (McCracken, Buckle et al., 2020; McCracken, Adams et al., 2020). In the case of growing bottles 0.4% (v/v) propanoic acid was also added. Cooked fly media was stored at 8°C for up to 2 weeks and was warmed before use. All experiments were done in a climate-controlled environment with a 12:12 hour light-dark cycle, at 25°C and ±60% humidity.

Experiments were conducted at two dietary conditions, 2% yeast (dietary restricted; DR) and yeast fully fed, with all other dietary components remaining the same. Where the GeneSwitch construct was used, food was supplemented with RU486 (Thermo Fisher Scientific) to induce transgene expression, or the same volume of absolute ethanol once the food cooled down (~80°C), as previously described in Hayman et al., 2025 (200 μM final media concentration; dissolved in 8.6 mL absolute ethanol (Fisher) per 1 L of fly media and mixed into the media) or control food lacking RU486 (but still containing 8.6 mL absolute ethanol per 1 L of fly media). Prior work by ourselves and others has not detected any effects of RU486 on longevity (Gautrey et al., 2025; Hayman et al., 2025; Ford et al., 2007; Biteau et al., 2010; Huang et al., 2014).

Where specified, 50 µl of 60 µM Borrelidin stock solution or a control (3% ethanol), was pipetted onto the experimental diet, for a final concentration of 3 µM Borrelidin, assuming even distribution within the top ~1 mL surface layer of food (Landis et al., 2020). To distinguish effects of borrelidin from its antimicrobial properties (Fang et al., 2015), we conducted a separate lifespan experiment where flies were treated with 50 µl of a broad-spectrum antibiotic (final concentration in the diet: 100 µg/ml ampicillin, 50 µg/ml vancomycin, 100 µg/ml neomycin and 100 µg/ml metronidazole), or equal volume of dH_2_O for 10 days we previously validated eradicates the microbiome (McCracken, Adams et al., 2020).

### Genotypes

All fly genotypes used in this study are listed in Table 3.

**Table 3.**
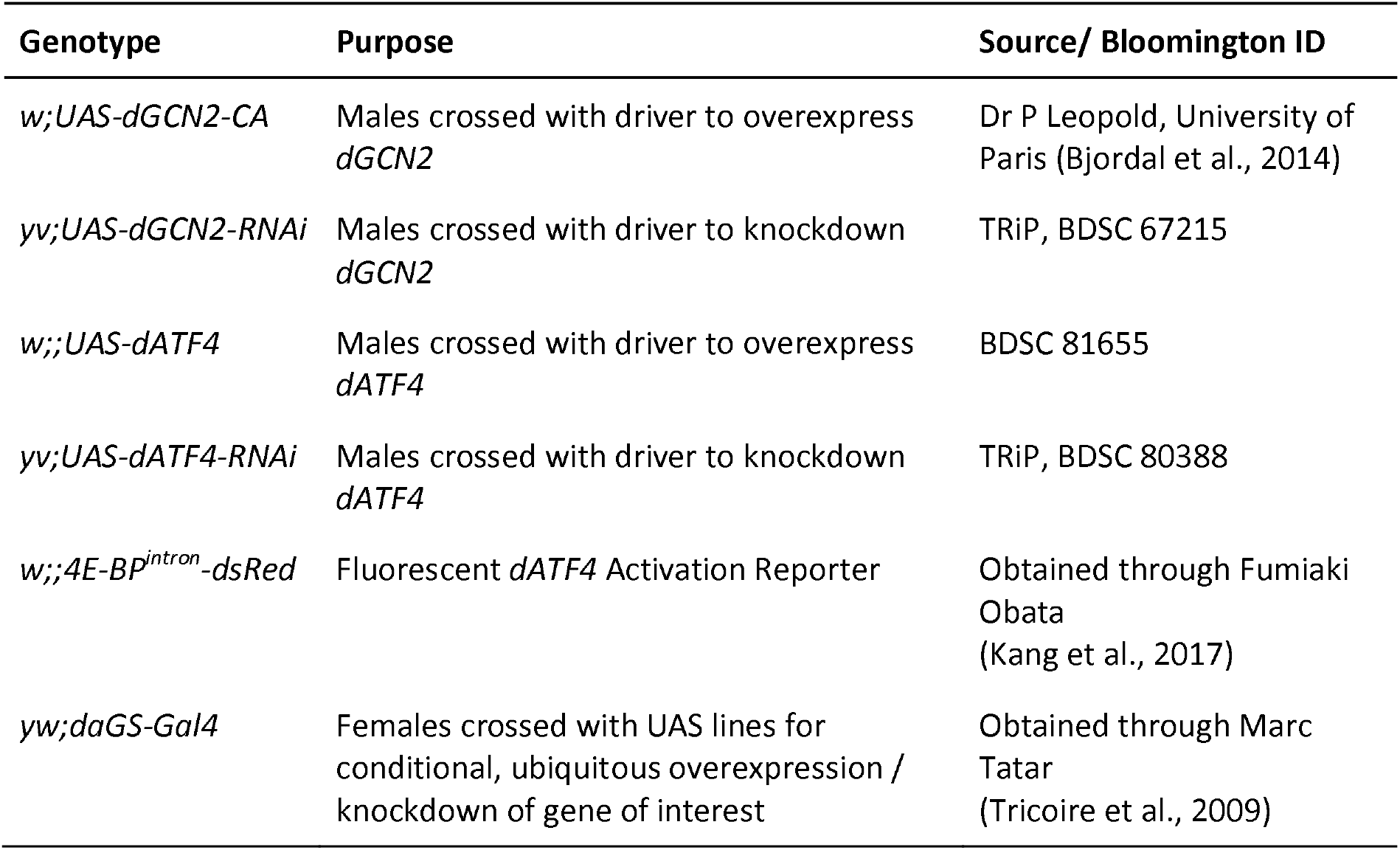
List of all fly genotypes used in this study and how they were obtained.

### Lifespan Experiments

All lifespan experiments were performed using females expressing the *daughterless* (*da*) GeneSwitch system (*daGS-GAL4*). The *da* promoter is active in almost all cells in the fly; using the GeneSwitch system therefore allows us to conditionally overexpress and knockdown (*in vivo* RNAi) genes in the whole fly (Tricoire et al., 2009). 3-5 males carrying various UAS transgenes (see Table 3) were crossed to 10 to 12 virgin females in bottles to control for density. As F1 progeny eclosed they were transferred to new bottles each day to generate age-matched cohorts, and left to mate for 2 days. Mated offspring were anaesthetised using carbon dioxide (Flystuff Flowbuddy; <5L / min), and females of the desired genotype were sorted into groups of up to 100 individuals and transferred to custom built demography cages (previously described in McCracken, Buckle et al., 2020; Good & Tatar, 2001). Flies were switched to experimental diets at age 3-4 days. After cageing mortality was measured every other day by conducting a fly census. At this time point, food vials were replaced, and dead flies were counted and removed from cages. Escaped and accidentally killed flies, and those stuck on the diet were right-censored (i.e., removed from mortality analysis as their cause of death could not be attributed to the experimental conditions tested).

### Imaging and quantification of dATF4 activity

A Leica M165 FC fluorescent dissection microscope was used to image flies expressing the *dATF4* reporter using dsRed filters, using LasX (Leica) image acquisition software. The following settings were used: exposure = 50ms, gain = 3.0, zoom = 1.25. Fluorescence intensity on images was quantified using Pixel Intensity Measurement in ImageJ. *dsRed* fluorescence of conditional *dATF4* overexpression and knockdown, was further quantified using fluorescent spectroscopy (*N* ≥ 10). Individual *dATF4* overexpression and *dATF4* knockdown flies were lysed with 5mm stainless steel beads (Qiagen) in PBS (1000µl PBS and 500µl PBS, respectively) for 3 minutes at 25 Hz using TissueLyser III (Qiagen). Samples were then centrifuged for 3 minutes at 13,000 rpm. The supernatant of samples was pipetted on 96-well, black-bottom cell culture microplates (Greiner BIO-ONE) and fluorescence was then measured using HIDEX Sense Beta Plus Microplate Reader (Type 425-311). The following settings were used: Technology = Fluorescence, Excitation = 560/40 nm, Emission = 590/20 nm, Mirror = Automatic, PMT voltage = 575 V, Flashes = 50 (*dATF4* knockdown) / 10 (*dATF4* overexpression). The volume of PBS and the number of flashes used to analyse *dATF4* knockdown flies was adjusted to account for the expected reduction in reporter activity and ensure measurements remain in the linear detection range of signal intensity. The average fluorescence of two technical replicates is given.

### Data Analysis for lifespan, fecundity, development and dATF4 activity

Data handling of lifespan data was done using a custom pipeline using spreadsheet software and R. Changes in lifespan associated with expression of different genes, dietary conditions and pharmacological interventions were analysed using the *coxme* package in R (https://CRAN.R-project.org/package=coxme). The models accounted for cage effects (using a random effect) and shared days of sorting into cages (batch) using a fixed effect. Each sorting day flies were split across different dietary conditions thereby balancing treatments. Changes in *dATF4* activity were analysed using ANOVA and post-hoc t-test to analyse imageJ results and Welch t-test for plate reader results.

#### RNA-sequencing library preparation

For RNA-seq analysis of *dATF4* overexpression and knockdown on both dietary conditions (DR and fully fed), were sampled from cages when 11 days old, after 10 days on RU486 or control diets. Samples were processed as a mix of 4-6 whole flies, snap frozen and kept at −70°C until analysis. Total RNA was extracted from 4-6 whole flies using Qiagen RNeasy Mini kits, following RNeasy Mini Kit Quick-Start Protocol. cDNA libraries for Oxford Nanopore sequencing were prepared using cDNA-PCR Barcoding Kit V14 SQK-PCB114.24 kit (Oxford Nanopore Technologies; ONT), following cDNA-PCR Sequencing V14 - Barcoding (SQK-PCB114.24) protocol with modifications. cDNA concentration was measured using Qubit and 1.925ng barcoded cDNA per sample was added to each library. In total three libraries were prepared to contain two biological replicates per condition to make up a total of 16 samples per flow cell. A total of three flow cells were run providing a N=6 per condition. Sequencing was done using GridION X5 (ONT).

#### Transcriptomic Analysis

For bioinformatic analysis we used minimap2 (v2.24; Li, 2018) and Salmon (v1.10.2; Patro et al., 2017) and we used the BDGP 6.32 genome and associated transcript annotation file as reference files. Prior to differential gene expression analysis, we filtered libraries with less than 400,000 reads. This resulted in exclusion of two libraries in the *dATF4* overexpression dataset (DR control and fully fed control) and three libraries in the *dATF4* knockdown dataset (2 DR control and 1 DR induced). Low abundance genes were filtered from the dataset to improve statistical power and minimise false positives. Genes were included in the analysis if they had a count-per-million (CPM) greater than 15 in all samples included in further analysis. The CPM threshold was selected based on exploratory data analysis to ensure reliable read-depth for reliable differential expression analysis, while maintaining sufficient gene representation. Differential expression analysis was done using the *edgeR* package and “glmFit” (Baldoni et al., 2024). Rank correlation between *dATF4* overexpression and *dATF4* knockdown was determined using Spearman’s rank correlation coefficient. Gene set enrichment analysis (GSEA; Korotkevich et al., 2021) was performed to identify biological pathways and processes associated with the differential gene expression profile. The KEGG (Kyoto Encyclopedia of Genes and Genomes; Kanehisa,2004) pathway database was used as reference gene sets for enrichment testing.

## Acknowledgements

MJPS is a Sir Henry Dale fellow (Wellcome and Royal Society: 216405/Z/19/Z). Dan Hayman is a Vivensa Foundation ECR Fellow (ECRF24\13). Further support from a Royal Society ISPF International Collaboration award to MJPS and FO (ICA\R2\242197). Stocks obtained from the Bloomington Drosophila Stock Center (NIH P40OD018537) were used in this study.

