## Supplementary material for "Suppression rather than activation of the integrated-stress-response (GCN2-ATF4) pathway extends lifespan in the fly"

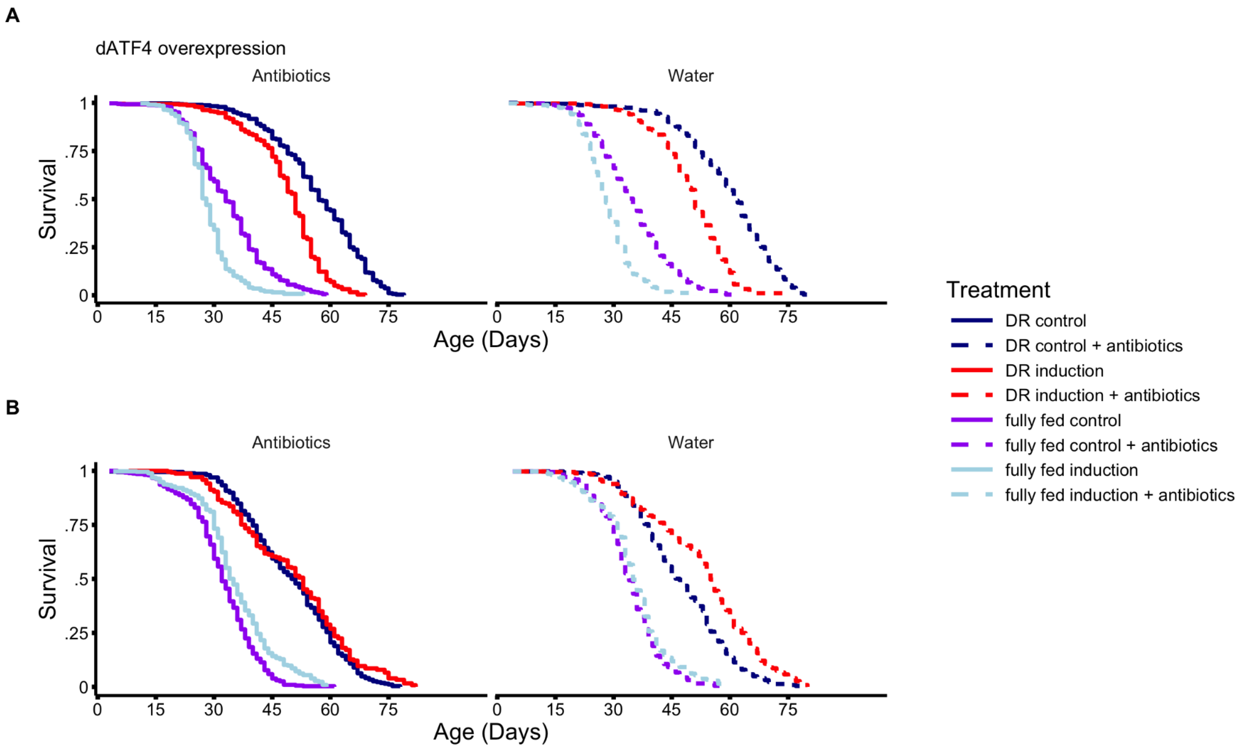

**Figure S1.** Microbial load does not affect the lifespan effect of *dATF4* manipulation. Survival curves of flies where the GeneSwitch construct was used to conditionally induce (A) overexpression or  (A) knockdown (*in vivo* RNAi) of *dATF4* to test the effect of microbial load on the lifespan effect of *dATF4* manipulation. Flies on antibiotic treatment were treated with 50µl of broadband antibiotics to eradicate their microbiome, control flies were treated with the same volume of $dH_2O$.  All sample sizes and statistics are given in Table S3.

11

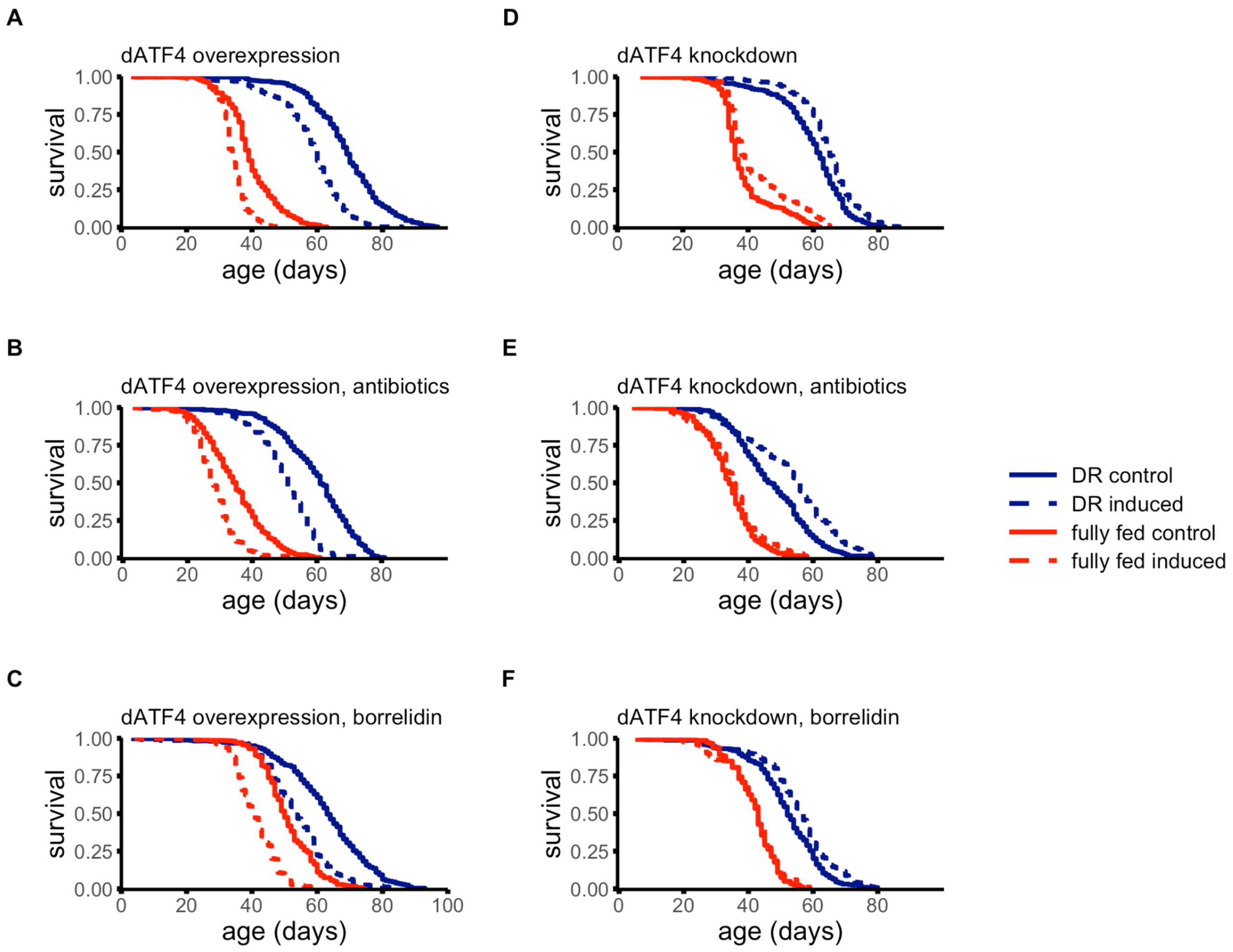

**Figure S2.** *dATF4* overexpression consistently reduces lifespan, while *dATF4* knockdown extends lifespan. Survival curves of flies where the GeneSwitch construct was used to conditionally induce (A-C) overexpression or (B-F) knockdown (*in vivo* RNAi) of *dATF4.* Replication was obtained from the control experiments for the “antibiotics” and “borrelidin” experiments and analysed separately. Note, these controls contained an additional 50µl $dH_2O$ or 3% ethanol respectively. All sample sizes and statistics are given in Table S4. Note, sample size of the knockdown fully fed group of dATF4 knockdown is low.

**Table S1. Statistics for the effect of genetic manipulation of the GCN2–ATF4 pathway on lifespan.** A) All sample sizes per treatment and condition. B-D) Statistics for effect of (B) conditional overexpression/knockdown of *dGCN2* and *dATF4*; (C) DR (2% yeast) in comparison to fully fed conditions (8% yeast); (D) the interaction of diet on gene induction on lifespan. B-D Statistics calculated using Coxme R package.

| A) | Sample Sizes | | | |
|---|---|---|---|---|
| **Condition** | **DR Induction** | **DR Control** | **fully fed Induction** | **fully fed Control** |
| *dGCN2* overexpression | 422 | 354 | 388 | 400 |
| *dGCN2* knockdown | 391 | 483 | 482 | 490 |
| *dATF4* overexpression | 441 | 376 | 487 | 487 |
| *dATF4* knockdown | 466 | 492 | 366 | 363 |
| B) | **Induction vs Control Statistics** | | | |
| **Condition** | **estimate** | **exp(estimate)** | **se** | **p-value** |
| *dGCN2* overexpression | 5.590 | 267.752 | 0.281 | < 0.0001 |
| *dGCN2* knockdown | -0.238 | 0.788 | 0.128 | 0.0628 |
| *dATF4* overexpression | 1.186 | 3.273 | 0.169 | < 0.0001 |
| *dATF4* knockdown | -0.549 | 0.577 | 0.120 | < 0.0001 |
| C) | **DR vs fully fed Statistics** | | | |
| *dGCN2* overexpression | -1.647 | 0.193 | 0.195 | < 0.0001 |
| *dGCN2* knockdown | -2.328 | 0.0975 | 0.138 | < 0.0001 |
| *dATF4* overexpression | -3.434 | 0.0323 | 0.169 | < 0.0001 |
| *dATF4* knockdown | -2.293 | 0.101 | 0.123 | < 0.0001 |
| D) | **Interaction Statistics** | | | |

| | | | | |
|---|---|---|---|---|
| *dGCN2* overexpression | 0.754 | 2.124 | 0.258 | < 0.01 |
| *dGCN2* knockdown | 0.311 | 1.365 | 0.187 | 0.0958 |
| *dATF4* overexpression | -0.00127 | 0.999 | 0.215 | 0.995 |
| *dATF4* knockdown | 0.07 | 1.082 | 0.161 | 0.624 |

**Table S2. Sample sizes and statistics for the effect of borrelidin treatment on *dATF4* manipulation.** A, B) All sample sizes per treatment in (A) *dATF4* overexpression and (B) dATF4 knockdown. C-D) Model following selection using backward removal of non-significant terms. (C) dATF4 overexpression and (D) dATF4 knockdown. B-D Statistics calculated using the *coxme* R package. Note sample sizes for fully fed (8% yeast) conditions are relatively low, and estimates are less certain for those treatments.

| A) | **Sample Sizes: *dATF4* overexpression** | | | |
|---|---|---|---|---|
| **Condition** | **DR Induction** | **DR Control** | **Fully fed Induction** | **Fully fed Control** |
| Control | 371 | 327 | 377 | 322 |
| Borrelidin | 304 | 386 | 249 | 271 |

| B) | **Sample Sizes: *dATF4* knockdown** | | | |
|---|---|---|---|---|
| **Condition** | **DR Induction** | **DR Control** | **Fully fed Induction** | **Fully fed Control** |
| Control | 336 | 326 | 161 | 147 |
| Borrelidin | 236 | 262 | 165 | 166 |

C) ***dATF4* overexpression**

| | **estimate** | **exp(estimate)** | **se** | **p-value** |
|---|---|---|---|---|
| Borrelidin | 0.329 | 1.39 | 0.148 | 0.038 |
| Induction | 1.91 | 6.75 | 0.220 | <0.0001 |
| DR | -1.35 | 0.260 | 0.238 | <0.0001 |
| Borrelidin:Induction | -0.950 | 0.387 | 0.232 | <0.0001 |
| Borrelidin:DR | 0.139 | 1.15 | 0.289 | 0.64 |
| Induction:DR | -1.02 | 0.361 | 0.327 | 0.002 |
| Borrelidin:Induction:DR | 0.843 | 2.32 | 0.417 | 0.04 |

**DR only**

| | | | | |
|---|---|---|---|---|
| Borrelidin | 0.470 | 1.60 | 0.259 | 0.07 |
| Induction | 0.852 | 2.34 | 0.270 | 0.002 |
| Borrelidin:Induction | -0.249 | 0.780 | 0.383 | 0.52 |
| **Fully fed only** | | | | |
| Borrelidin | 0.349 | 1.417 | 0.170 | 0.04 |
| Induction | 2.31 | 10.1 | 0.290 | <0.0001 |
| Borrelidin:Induction | -1.25 | 0.288 | 0.251 | <0.0001 |

D) ***dATF4* knockdown**

| | estimate | exp(estimate) | se | p-value |
|---|---|---|---|---|
| Borrelidin | -0.093 | 0.911 | 0.145 | 0.52 |
| DR | -1.32 | 0.266 | 0.142 | <0.0001 |
| Induction | -0.214 | 0.807 | 0.079 | <0.01 |
| Borrelidin:DR | 0.584 | 1.794 | 0.181 | 0.001 |
| **DR only** | | | | |
| Borrelidin | 0.348 | 1.42 | 0.099 | <0.001 |
| Induction | -0.319 | 0.727 | 0.087 | <0.001 |

| | | | | |
|---|---|---|---|---|
| Borrelidin:Induction | 0.265 | 1.30 | 0.131 | 0.04 |
| **Fully fed only** | | | | |
| Borrelidin | 0.067 | 1.07 | 0.308 | 0.83 |
| Induction | -0.059 | 0.943 | 0.277 | 0.83 |
| Borrelidin:Induction | -0.391 | 0.676 | 0.391 | 0.32 |

**Table S3. Sample sizes and statistics for the effect of antibiotic treatment on *dATF4* manipulation.** A, B) All sample sizes per treatment in (A) *dATF4* overexpression and (B) *dATF4* knockdown. C-D) Statistics for the effect of antibiotics in (C) *dATF4* overexpression and (D) *dATF4* knockdown. B-D Statistics calculated using the *coxme* R package.

| A) | **Sample Sizes: *dATF4* overexpression** | | | |
|---|---|---|---|---|
| **Condition** | **DR Induction** | **DR Control** | **Fully fed Induction** | **Fully fed Control** |
| Control | 322 | 434 | 429 | 427 |
| Antibiotics | 458 | 478 | 340 | 448 |

| B) | **Sample Sizes: *dATF4* knockdown** | | | |
|---|---|---|---|---|
| **Condition** | **DR Induction** | **DR Control** | **Fully fed Induction** | **Fully fed Control** |
| Control | 347 | 356 | 358 | 349 |
| Antibiotics | 241 | 360 | 443 | 484 |

C) ***dATF4* overexpression**

| | **estimate** | **exp(estimate)** | **se** | **p-value** |
|---|---|---|---|---|
| Antibiotics | 0.129 | 1.138 | 0.132 | 0.330 |
| RU | 1.128 | 3.091 | 0.104 | < 0.0001 |
| Antibiotics:RU | -0.074 | 0.928 | 0.194 | 0.702 |

D) ***dATF4* knockdown**

| | **estimate** | **exp(estimate)** | **se** | **p-value** |
|---|---|---|---|---|
| Antibiotics | 0.012 | 1.012 | 0.106 | 0.913 |
| RU | -0.364 | 0.695 | 0.109 | < 0.001 |
| Antibiotics:RU | 0.023 | 1.023 | 0.1522 | 0.880 |

**Table S4. Sample sizes and statistics for dATF4 overexpression and knockdown replicates.** A, B) All sample sizes per replicate in (A) *dATF4* overexpression and (B) *dATF4* knockdown. C,D) We analysed using coxme the effect of the induction of the transgene for each replicate, whilst controlling for diet (not shown). The main effect and the interaction with diet are reported. (C) *dATF4* overexpression and (D) *dATF4* knockdown.

A) **Sample Sizes: *dATF4* overexpression**

| **Condition** | **DR Induction** | **DR Control** | **8% Induction** | **8% Control** |
|---|---|---|---|---|
| Main | 441 | 376 | 487 | 487 |
| Antibiotics, control | 322 | 434 | 429 | 427 |
| Borrelidin control | 371 | 367 | 377 | 322 |

B) **Sample Sizes: *dATF4* knockdown**

| **Condition** | **DR Induction** | **DR Control** | **8% Induction** | **8% Control** |
|---|---|---|---|---|
| Control | 466 | 492 | 366 | 363 |
| Antibiotics, control | 347 | 356 | 358 | 349 |
| Borrelidin, control | 336 | 326 | 161 | 147 |

C) ***dATF4* overexpression**

| | **estimate** | **exp(estimate)** | **se** | **p-value** |
|---|---|---|---|---|
| Main, Induction | 1.186 | 3.273 | 0.169 | < 0.0001 |
| Main, Induction:DR | -0.00127 | 0.999 | 0.215 | 0.995 |
| Antibiotics, control, Induction | 1.101 | 3.008 | 0.186 | < 0.0001 |
| Antibiotics, control, Induction:DR | 0.095 | 1.100 | 0.272 | 0.727 |
| Borrelidin, control, Induction | 1.862 | 6.441 | 0.205 | < 0.0001 |
| Borrelidin, control, Induction:DR | -0.844 | 0.420 | 0.285 | < 0.01 |

D) ***dATF4* knockdown**

| | estimate | exp(estimate) | se | p-value |
|---|---|---|---|---|
| Main, Induction | -0.549 | 0.577 | 0.120 | < 0.0001 |
| Main, Induction:DR | 0.07 | 1.082 | 0.161 | 0.624 |
| Antibiotics, control, Induction | -0.203 | 0.816 | 0.104 | < 0.05 |
| Antibiotics, control, Induction:DR | -0.355 | 0.701 | 0.140 | < 0.05 |
| Borrelidin, control, Induction | -0.247 | 0.781 | 0.119 | < 0.05 |
| Borrelidin, control, Induction:DR | -0.300 | 0.741 | 0.242 | 0.215 |

**Table S5. Genes that changed significantly (p < 0.01) in expression and in opposite directions across both conditions (*dATF4* overexpression and knockdown).** Annotation and the nearest human paralog was collated from Flybase. When prior literature appeared to link the gene to ATF4, this reference is listed and a short conclusion is provided. Literature search was conducted with the gene name and ATF4 and a quick manual scan of the associated literature in both Google and Google Scholar.

| | *dATF4* overexpression | | | | *dATF4* knockdown | | | | | | |
|---|---|---|---|---|---|---|---|---|---|---|---|
| **Fly Base ID** | **logFC** | **PValue** | **FDR** | **symbol** | **logFC** | **PValue** | **FDR** | **annotation** | **human paralog** | **reference** | **conclusion** |
| FBgn01976 | -0.47 | 0.001 | 0.023 | Psf2 | 0.37 | 0.009 | 0.138 | initiation of DNA replication | GINS2 | | |
| FBgn0261524 | -0.37 | 0.009 | 0.089 | lic | 0.41 | 0.005 | 0.110 | MAP kinase kinase | MAP2K3 | | |
| FBgn0022224 | -0.28 | 0.009 | 0.089 | ubl | 0.34 | 0.002 | 0.079 | Ubiquitin-like | UBL5 | Wang et al., 2023 | thought to be independent death pathway of ATF4-CHOP |
| FBgn0035805 | 0.25 | 0.010 | 0.094 | CG7506 | -0.33 | 0.002 | 0.084 | participates in the c-ring assembly of mitochondrial ATP synthase | TMEM70 | | |
| FBgn0005278 | 0.31 | 0.010 | 0.094 | Sam-S | -0.33 | 0.010 | 0.142 | S-adenosylmethionine synthetase | MAT2A | Chen et al., 2021 | appears known ATF4 target |
| FBgn0029755 | 0.32 | 0.004 | 0.060 | Sas10 | -0.41 | 0.001 | 0.078 | Ribosomal biogenesis | UTP3 | | No direct connection but ribosomal biogenesis fits longevity effect |
| FBgn0015221 | 0.34 | 0.009 | 0.089 | Fer2LCH | -0.37 | 0.005 | 0.111 | Iron storage, one of the two subunits of insect ferritin | | Tang et al., 2024 | ATF4 regulates ferroptosis |
| FBgn0028473 | 0.48 | 0.002 | 0.042 | Non1 | -0.44 | 0.006 | 0.119 | GTP binding protein | GTPBP4 | | |
| FBgn0032144 | 0.55 | 0.007 | 0.078 | CG17633 | -0.93 | < 0.0001 | 0.005 | Proteolysis, metallocarboxypeptidase activity | CPA1 | | |
| FBgn0035666 | 0.63 | 0.007 | 0.078 | Jon65Aii | -0.64 | 0.007 | 0.127 | Proteolysis and immune system | | | |
| FBgn0034871 | 0.70 | 0.002 | 0.040 | CG3906 | -0.73 | 0.002 | 0.079 | unknown | | | |
| FBgn0036007 | 0.99 | 0.000 | < 0.0001 | path | -0.38 | 0.009 | 0.137 | amino-acid transporter | SLC36A1 | Zhang et al., 2018 | ATF4 regulates amino acid transporters and this likely include SLC36A1 |
| FBgn0033446 | 1.25 | 0.000 | 0.008 | CG1648 | -1.01 | 0.002 | 0.081 | unknown | | | |
